# Assessing the relation between protein phosphorylation, AlphaFold3 models and conformational variability

**DOI:** 10.1101/2025.04.14.648669

**Authors:** Pathmanaban Ramasamy, Jasper Zuallaert, Lennart Martens, Wim F. Vranken

**Affiliations:** CompOmics, VIB Center for Medical Biotechnology, VIB, Ghent, Belgium; Department of Medical Protein Research, Faculty of Medicine and Health Sciences, Ghent University, Ghent, Belgium; Interuniversity Institute of Bioinformatics in Brussels, ULB-VUB, Brussels, Belgium; Structural Biology Brussels, Vrije Universiteit Brussel, Brussels, Belgium; BioOrganic Mass Spectrometry Laboratory (LSMBO), IPHC UMR 7178, University of Strasbourg, CNRS, ProFI FR2048, Strasbourg, France

**Keywords:** Protein structures, conformational diversity, AlphaFold3, post-translational modifications (PTMs), phosphorylation

## Abstract

Proteins perform diverse functions critical to cellular processes. Transitions between functional states are often regulated by post-translational modifications (PTMs) such as phosphorylation, which dynamically influence protein structure, function, folding, and interactions. Dysregulation of PTMs can therefore contribute to diseases such as cancer and Alzheimer’s. However, the structure-function relationship between proteins and their modifications remains poorly understood due to a lack of experimental structural data, the inherent diversity of PTMs, and the dynamic nature of proteins. Recent advances in deep learning, particularly AlphaFold, have transformed protein structure prediction with near-experimental accuracy. However, it remains unclear whether these models can effectively capture PTM-driven conformational changes, such as those induced by phosphorylation. Here, we systematically evaluated AlphaFold models (AF2, AF3-non phospho, and AF3-phospho) to assess their ability to predict phosphorylation-induced structural diversity. By analysing experimentally derived conformational ensembles, we found that all models predominantly aligned with dominant structural states, often failing to capture phosphorylation-specific conformations. Despite its phosphorylation-aware design, AF3-phospho predictions provided only modest improvement over AF2 and AF3-non phospho predictions. Our findings highlight key challenges in modelling PTM-driven structural landscapes and underscore the need for more adaptable structure prediction frameworks capable of capturing modification-induced conformational variability.

## 1 Introduction

Proteins are essential biomolecules that play a myriad of roles in cellular processes, serving as enzymes, structural components, signaling molecules, and more. Their function is intricately linked to their three-dimensional structure, which is determined by the sequence of amino acids and stabilized by various interactions such as hydrogen bonds, disulfide bridges, hydrophobic effects, and ionic interactions^1–3^. This precise structure-function relationship is fundamental for the protein’s biological activity^4^. Post-translational modifications (PTMs), including phosphorylation, ubiquitination, glycosylation, and methylation, further diversify protein function and regulation^5^. PTMs are critical in modulating protein activity, stability, localization, and interaction with other molecules, thereby fine-tuning cellular processes and responses^5–7^. Among these, phosphorylation is one of the most extensively studied due to its pivotal role in regulating cellular processes^7,8^, such as the MAPK/ERK pathway, where sequential phosphorylation events propagate signals that control cell growth, differentiation, and apoptosis^9^. Dysregulation of phosphorylation is implicated in numerous diseases, including cancer, diabetes, and neurodegenerative disorders^10,11^. Phosphorylation can induce significant conformational changes in the protein, acting as a molecular switch that regulates protein activity^5,6^.

Understanding the mechanisms of PTMs, particularly phosphorylation, is crucial for fully comprehending protein function and, by extension, for the development of targeted therapies. Though 60% of the human proteome is known to be phosphorylated^8^, only a small fraction of these known phosphosites are associated with functional relevance^12^. Computational and experimental approaches have tried to identify such functionally important phosphosites^13,14^. However, these methods cannot identify the structural impact of phosphorylation, which is pivotal in how signal transduction, enzyme regulation, and other cellular processes^6,15,16^ are mediated. Our incomplete understanding of how phosphorylation modulates protein conformation especially impacts analysis of large-scale proteomics datasets, given the dearth of experimentally available structural information^5,8^.

AlphaFold2, a deep learning-based structure prediction algorithm, has demonstrated remarkable performance in protein structure prediction from amino acid sequence^17^. This tool has been instrumental in elucidating the structures of many previously unexplored or challenging proteins, resulting in for example a dramatic increase of the structural coverage of the human proteome and interactome^18–20^. AlphaFold2 excels at predicting static ground-state structures but often misses dynamic changes from modifications like phosphorylation^21–23^. AlphaFold3 improves on this by modelling ligands, complexes, and modifications, offering better insight into functional protein states^24^.

However, proteins are not static entities; they exist as ensembles of interchanging conformational states^22^. This conformational diversity, especially in phosphorylated proteins, presents a challenge for AlphaFold and similar computational models, which do not capture gradations in dynamical properties^25^. Trained predominantly on experimental structures from the Protein Data Bank (PDB) determined by x-ray diffraction, with the protein in crystallized form at typically cryogenic temperatures, AlphaFold essentially excels at predicting the most stable conformations in these conditions. How well AlphaFold can sample multiple protein conformations beyond these dominant structural states remains unclear^26,27^; its ability to model conformational diversity is not fully understood and was only validated on a limited number of well-studied experimentally determined cases. Its predictions may therefore be guided more by pattern recognition for the most probable conformer than by an intrinsic understanding of the energetics underlying protein ensembles^27,28^.

These open questions on AlphaFold’s performance on highly dynamic proteins are ^27,28^particularly relevant for phosphorylation, which is known to drive distinct structural states critical for regulatory functions. To date, no large-scale studies have systematically assessed AlphaFold’s performance on phosphorylated proteins in light of their complex conformational landscapes. The distinct structural states introduced by phosphorylation are often functionally critical, yet transient or less abundant - can AlphaFold accurately model such structural changes, or does it merely default to predicting the dominant, non-phosphorylated conformation?

Here, we provide a large-scale, detailed assessment of the conformational diversity of phosphorylated proteins by analyzing multiple experimentally observed conformational states of eukaryotic proteins (human, mouse, and rat) in the Protein Data Bank (PDB)^29^. In addition, we evaluate the ability of AlphaFold models (AF2, AF3) to predict these phosphorylation-induced conformational states, focusing on both global and local structural changes. While AF2/3 was trained on PDB structures with limited conformational variability, here we use the same structural information to evaluate model predictions in a distinct context, focusing on conformational diversity and structural transitions rather than direct replication of training data. Using metrics such as Conformational Diversity Score (CDS), Phospho-Adjusted CDS, co-evolutionary signals and clustering approaches, we analyze how phosphorylation drives conformational shifts and assess whether AlphaFold models (AF2, AF3) capture these transitions. Our analysis highlights both the strengths and limitations of AlphaFold (AF2, AF3) in modelling phosphorylated proteins, revealing biases toward dominant structural ensembles and reduced sensitivity to phosphorylation-induced conformational changes. Notably, we observed that AF models prioritize evolutionarily stable conformations, favouring those reinforced by co-evolutionary constraints rather than modelling the full spectrum of structural variability. Our findings provide a foundation for improving predictive algorithms by emphasizing the need to capture local structural variations and rare conformations. Future refinements of AlphaFold architectures-such as incorporating context-specific embeddings or hybrid frameworks-may enable more accurate modelling of post-translationally modified protein states.

## 2 Methods

### 2.1 Phosphoprotein structure selection and curation

We obtained all available PDB structures, chains, and modeled segments for human, mouse and rat canonical proteins from UniProtKB^30^. For each protein we retrieved all available protein structure information from PDB^29^. The modified amino acid information was obtained to get the phosphorylated structures and their non-phospho counterparts (where the same residue is phosphorylated in one and not phosphorylated in other) by querying the PDB database. To identify structures crystallized with phosphorylation modifications, we conducted a comprehensive API screening of the PDB repository (https://data.rcsb.org/#rest-api) (19 September 2024) and retrieved modified residue information for all structures per protein. We selected phosphoproteins crystallized with one or more canonical phosphosites: phosphoserine (SEP), phosphothreonine (TPO), and phosphotyrosine (PTR). For the non-phospho counterparts, we ensured the phospho and non-phospho PDB IDs are mapped to the same UniProt ID, and the residues are not phosphorylated (SER, THR and TYR). This resulted in 185 proteins covering a total of 2015 phosphosites from 1327 phospho structures and 3867 non-phospho structures. All the PDB files were downloaded on 20 September 2024. To ensure the accuracy of conformational details and the completeness of modelled segments, we included only structures where the modelled fragments of the protein contained at least 50 amino acids.

For each modified amino acid in the structure, we assessed missing residues and prioritized structures with no missing residues. If no such structures were available, we minimized the number of missing residues around both phosphorylated (P data) and non-phosphorylated sites (NP data). For phosphorylated structures, we selected one representative structure per modification state (single, dual, or multi-phosphorylated) based on minimal missing residues and with better resolution. For non-phosphorylated structures, all entries meeting the filtering criteria were retained. This approach ensured the inclusion of PDB structures covering sufficiently large protein fragments with complete information around the modified/unmodified residues. The final filtered data resulted in 109 different proteins covering a total of 253 phosphosites from 159 phospho structures and 1695 non-phospho structures.

To cover diverse phosphosite and conformational preferences, proteins were selected such that the number of phosphosites in a structure ranged from one to five, encompassing a combination of SEP, TPO, and PTR. Details of all phosphoproteins, including the number of PDB structures modeled with and without phosphorylation and phosphosite information are presented in Table 1.

### 2.2 Protein conformers and conformational variability in protein structures

We assessed the conformational diversity of proteins in the context of post-translational modifications, with a specific focus on phosphorylation. We filtered protein chains according to the criteria previously outlined (Section 2.1) and labelled each chain as P (phosphorylated) or NP (non-phosphorylated) based on its phosphorylation status. This allowed us to quantify and compare the number of P and NP conformers available for each protein, providing insights into the structural variability associated with phosphorylation.

To investigate conformational diversity between P and NP protein conformers, we categorized structural changes by assessing pairwise Root Mean Square Deviation (RMSD) values. Both global and segment-based RMSD values were analyzed to capture variations at different structural scales. We performed structural alignments using the PDB Superimposer module in BioPython^31^. For global alignment, all C-alpha atoms common to both structures were used to calculate the root mean square deviation (RMSD). For segment alignment, only the residues within a defined region (e.g., ±15 residues around phospho-sites) were included. The alignment utilized the Kabsch algorithm to determine the optimal rotation and translation matrices that minimized the RMSD between the selected atoms. This approach ensures sensitivity to both global conformational changes and localized structural deviations.

In order to determine the distinct conformational state of a protein, we clustered all P and NP conformers for a specific protein based on pairwise structural alignment. We performed both the global alignment (on full chain) and segment alignment (spanning 15 amino acids upstream and downstream in sequence space) using PDB Superpose method. The RMSD values obtained between each pair were used for clustering. Hierarchical clustering was performed using the average linkage method, with clusters defined based on a dynamic distance threshold set at 70% of the maximum linkage distance to ensure appropriate separation of structural conformations. A dendrogram was generated to visualize the clustering results, with distinct branch colours indicating major structural clusters and the cluster threshold visually marked to highlight structural variation patterns. Each cluster therefore represents a distinct conformational state of the protein. We considered the number of clusters per protein as a quantitative measure of conformational diversity. To assess the influence of phosphorylation on conformational diversity, we annotated each cluster with the number of P and NP conformers. This allowed us to explore whether phosphorylation affects conformational states by stabilizing or favouring specific clusters.

To further quantify conformational diversity, we calculated a Conformational Diversity Score (CDS) and an adjusted phospho Conformational Diversity Score (pCDS). The CDS provided a baseline measure of structural variability, while the adjusted pCDS accounted for the relative proportions of P and NP conformers, normalizing the score to evaluate the specific contribution of phosphorylation to structural diversity.

#### 2.2.1 Conformational Diversity Score (*cds*)

The Conformational Diversity Score (*cds*) is used to quantify the structural diversity of protein conformations within a set of structural clusters. The CDS considers both the number of clusters a protein adopts and the relative size of each cluster, thus providing an overall measure of conformational flexibility.

The CDS for a protein is calculated as follows:

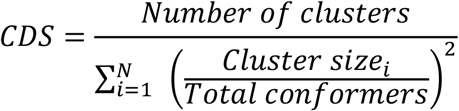

where:

- *Number of clusters*, the total number of unique structural clusters identified for a given protein.
- *N*, the total number of clusters.
- *Cluster size*_*i*_, the number of structures in cluster.
- *Total conformers*, the sum of all conformers across all clusters for the protein.

This formula captures the spread of conformers across clusters, with higher CDS values indicating more even distribution across clusters (greater diversity), and lower CDS values suggesting concentration within fewer clusters (less diversity).

#### 2.2.2 Adjusted phospho-CDS (pCDS)

To investigate the impact of phosphorylation on conformational diversity, an adjusted phospho-CDS (*pCDS*) was introduced. This score incorporates information about the distribution of P and NP conformers within each cluster, assigning higher weight to clusters where both phosphorylation (P and NP) states are present in a balanced manner. *pCDS* aims to highlight clusters where phosphorylation induces distinct conformational states, while still accounting for structural diversity across the dataset.

The adjusted phospho-CDS is calculated as:

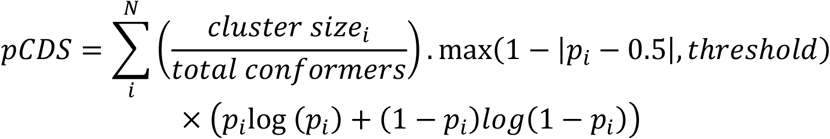

Where:

- *p*_*i*_ the proportion of phospho structures within cluster *i*, calculated as 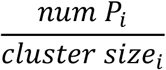
- (1 − |*p*_*i*_ − 0.5|, *threshold*), serves as a weighting term that increases when clusters have balanced P and NP structures, emphasizing clusters where phosphorylation may influence conformational shifts.
- *threshold* is set as a minimum value (e.g., 0.3) to ensure that clusters with high P or NP dominance still contribute meaningfully to the score.

This formula captures the diversity upon phosphorylation by emphasizing clusters that exhibit mixed modification states, providing a nuanced view of structural adaptability in response to phosphorylation.

### 2.3 Domain regions, secondary structural states and solvent accessibility

Domain information for the proteins was obtained using the InterPro^32^ API, which provides annotations for protein domains and their coordinates within UniProt sequences. The domain regions were then mapped onto modelled PDB structures, the PDBe API was used to retrieve residue ranges for specific PDB chains. The InterPro API outputs were filtered to include only domain annotations, while the PDBe API results were cross-referenced to match PDB IDs and chains corresponding to the modelled structures. These data were integrated to identify and compare domain regions across P and NP conformers. Secondary structural information for proteins was derived using STRIDE^33^, which annotates eight secondary structure classes based on PDB files. These classes were subsequently mapped into three categories: helix, strand, and coil. Relative solvent accessibility (RSA) was calculated as a percentage of their maximum solvent-accessible area, using established values for standard amino acids^33,34^. STRIDE was run on modified PDB files, ensuring phosphorylated residues were correctly processed, and the results were matched to specific residues in both P and NP states.

### 2.4 Protein structures (PDB, AlphaFold2, AlphaFold3 predictions)

All PDB structures used in this analysis were obtained from the RCSB PDB (https://www.rcsb.org/) on 20 September 2024. AlphaFold2 structures were sourced from the AlphaFold2 Database (https://alphafold.ebi.ac.uk/) on 08 October 2024 and were labelled AF2.

AlphaFold3 structures were predicted using the AlphaFold3 server (https://alphafoldserver.com/). A single Json file was used for batch predictions as described in the server (https://github.com/google-deepmind/alphafold/blob/main/server/README.md). AF3 prediction jobs were submitted between 02 October 2024 and 08 October 2024.

#### 2.4.1 AlphaFold3 predictions

##### Unphosphorylated structure predictions

AlphaFold3 predictions were generated from the corresponding canonical UniProtKB protein sequences without any modifications. These predictions were labelled as AF3.

##### Phosphorylated structure predictions

Due to the server limitations of 20 structures per day, we only predicted the phosphorylation status of the conformer that showed the maximum conformational variability in a protein. These predictions were labelled as AF3-p.

We used the MAXIT suite (https://sw-tools.rcsb.org/apps/MAXIT/index.html) to convert the PDBx/mmCIF format to PDB format.

### 2.5 Structure comparison and evaluation metrics

To evaluate the performance of predicted models; AlphaFold2 (AF2), AlphaFold3 (AF3), and AlphaFold3 phosphorylation-specific (AF3-p) various metrics and comparison strategies were employed. Experimental Protein Data Bank (PDB) structures, including both phosphorylated and non-phosphorylated forms, were used to cross-validate the predictions. Key validation metrics included the consistency of Root Mean Square Deviation (RMSD) values, structural integrity, and conformational accuracy, particularly at phosphorylation sites and surrounding residues.

To analyze local conformational changes around phosphorylation sites, residue contacts of phosphorylated residues (in P conformers) and their corresponding non-phosphorylated residues (in wild-type NP conformers) were measured within a 8Å radius^35–37^.

### 2.6 Hierarchical clustering and heatmap visualization

To evaluate the structural alignment of AF2, AF3, and AF3-p predictions with experimental P) and NP conformations, hierarchical clustering was performed using distance matrices derived from three metrics:

1. **global RMSD**: Root mean square deviation calculated across the entire protein structure.
2. **segment RMSD**: RMSD values calculated for the ±15-residue region surrounding phosphorylation sites (N- and C-terminal extensions).
3. **contact map differences**: Residue contact differences within a 8Å radius^35–37^ around P and NP sites.

For each protein, representative structures were chosen for each distinct phosphorylation state from PDB entries as described in section 2.1. Predictions from AF2, AF3, and AF3-p were aligned to these representative P and NP conformers to calculate pairwise distances, which were used to generate condensed distance matrices. Hierarchical clustering was then performed using the average linkage method, and the resulting dendrograms were visualized as heatmaps, where clustering patterns were used to evaluate the model’s proximity to structural ensembles.

Heatmaps were generated to illustrate the relative RMSD and contact differences for selected proteins, highlighting how model predictions align with experimentally observed conformers. These visualizations provide insights into the structural tendencies of each model, particularly in capturing phosphorylation-related conformations.

### 2.7 Residue Interaction Networks (RINs) and co-evolutionary analysis

To evaluate how AF models capture phosphorylated (P) and non-phosphorylated (NP) conformers, we constructed Residue Interaction Networks (RINs) and inferred co-evolutionary constraints from EvCouplings^38^. Global RINs were built from PDB structures, where residue-residue contacts were defined within 8.0 Å, including phosphorylated residues (SEP, TPO, PTR). Localized subgraphs captured P and NP residues along with their immediate neighbours. Network properties (degree centrality, edge betweenness, clustering coefficient, and eigenvector centrality) were computed to assess residue contributions to structural stability. Co-evolving residue pairs were inferred from EvCouplings and mapped onto RINs as weighted edges based on evolutionary coupling probabilities. The analysis was performed for global structures and localized subgraphs, capturing both large-scale and site-specific constraints. AF model predictions were compared to PDB conformers using Jaccard similarity, assessing overlap in residue interaction and co-evolutionary networks:

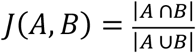

where A and B represent residue contacts in AF and PDB networks. Mean similarity scores were computed for global RINs and subgraphs to quantify model alignment with PDB conformers.

### 2.8 Statistical test for significance

Statistical analyses were performed to assess the effect of ligand binding on conformational diversity, a Tukey Honest Significant Difference (HSD) test was performed separately for P-P, P-NP, and NP-NP pairs. RMSD distributions across the three ligand categories (ligand-bound, mixed, no-ligand) were compared within each pair type. Statistical significance was determined based on an adjusted significance threshold (α=0.05) to account for multiple comparisons. To evaluate whether ligand binding altered the structural relationship between P and NP conformers, Pearson correlation coefficients were calculated between RMSD values of P-P, P-NP, and NP-NP pairs within each ligand category. Correlation strength was assessed using the pearson correlation coefficient (r), and statistical significance was determined by the corresponding p-value. Correlation analyses were conducted only for pairs where RMSD values were available for both pair types.

To evaluate the performance of AF2, AF3, and AF3-p in predicting phosphorylated (P) and non-phosphorylated (NP) conformations, as well as their structural and label alignment tendencies, chi-square test and Kruskal-Wallis test was performed. To determine whether the proportions of predictions aligning with P versus NP conformers differed significantly across models, a chi-square test of independence was applied, testing for overall distribution differences.

The Kruskal-Wallis test was used to compare the proportions of predictions aligning with the same cluster as the closest conformer, providing a non-parametric measure of structural consistency across the models. For label alignment (P vs. NP), binomial tests were conducted to assess whether the proportion of “same label” matches for each model exceeded a random baseline (p=0.5), highlighting each model’s ability to differentiate between phosphorylated and non-phosphorylated states. Additionally, a chi-square test was performed to compare the overall distribution of “Same Label” and “different label” cases across models to assess inter-model differences in label alignment performance.

All statistical analyses were conducted using Python, primarily utilizing the scipy library. A p-value threshold of 0.05 was used to determine statistical significance, ensuring robustness in identifying meaningful differences among the models.

## 3 Results

We retrieved all available protein structures from the Protein Data Bank (PDB) for human, mouse, and rat to investigate conformational changes upon phosphorylation. Of the 10,686 proteins with at least one structure in the PDB, only 8.5% had at least one phosphorylated canonical site, identified by residue codes for phosphoserine (SEP), phosphothreonine (TPO), or phosphotyrosine (PTR). Filtering for phosphorylated structures with ≥50 amino acids reduced this to 2.2%, indicating that most phospho-containing entries are short peptides. When restricting to proteins with both phosphorylated and non-phosphorylated structures, the number dropped further to 1%, underscoring the limited availability of full-length phosphorylated proteins (Figure 1A). Most phosphorylated structures belong to the kinase superfamily, followed by small GTPases and heat shock proteins (Figure 1B). Singly phosphorylated proteins are most common, with TPO being the most frequent modification, followed by PTR and SEP. For dual phosphorylation, PTR-PTR combinations dominate, followed by SEP-TPO pairs.

**Figure 1:**
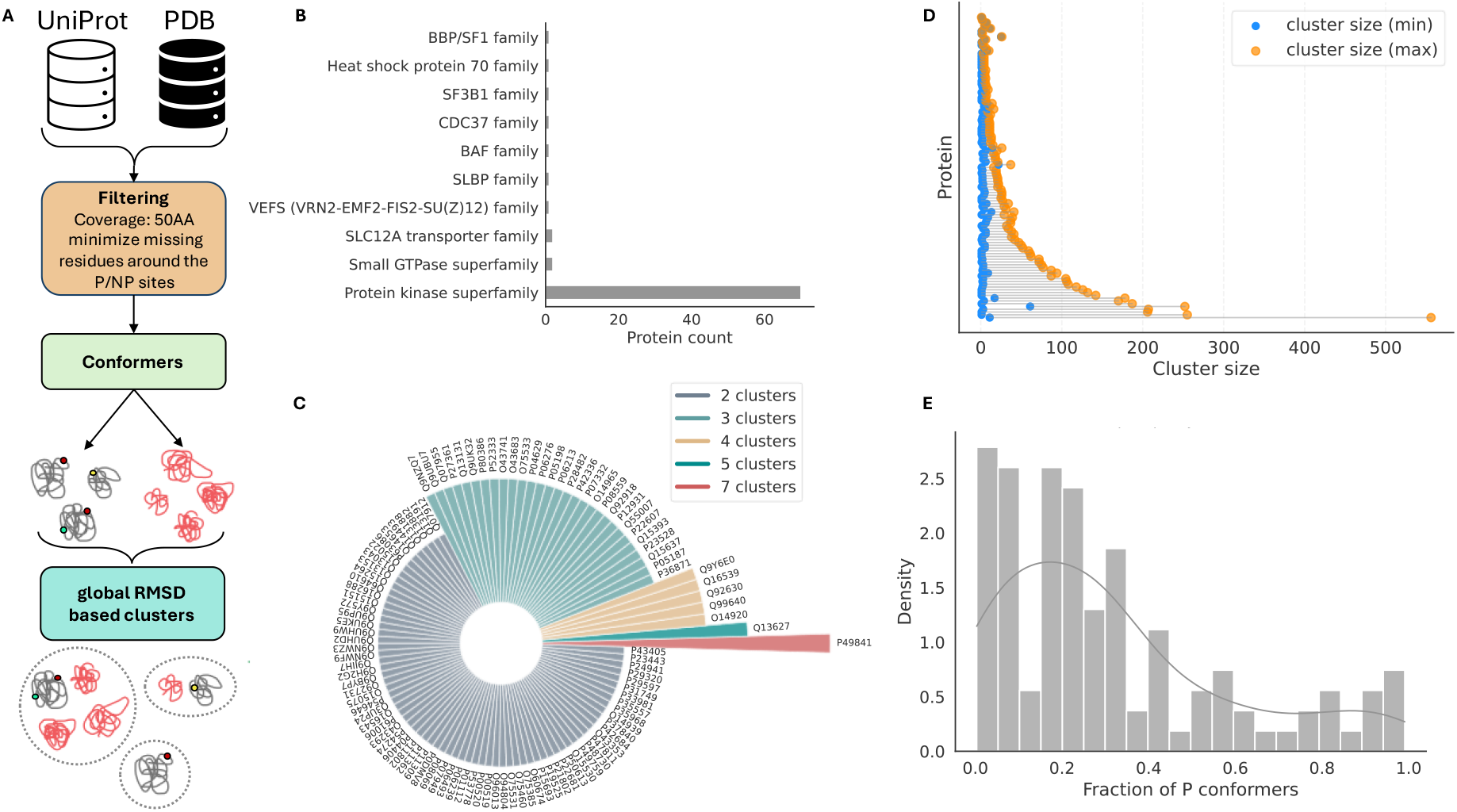
Overview of phosphorylated protein structures in the PDB. **A)** Workflow for collecting, filtering, and clustering phosphorylated (P) and non-phosphorylated (NP) protein conformers. The process involves retrieving structures from UniProt and PDB, filtering for complete residues surrounding phosphosites, and clustering based on global RMSD values. **B)** Protein family distribution of experimentally determined phosphorylated structures in the PDB, dominated by kinase superfamilies, with smaller contributions from other families. **C)** Sunburst plot showing the number of distinct clusters per protein based on global RMSD clustering, with proteins classified by cluster count. **D)** Dumbbell plot illustrating cluster size statistics for each protein. The x-axis represents the number of conformers per cluster, with blue and orange markers indicating the smallest and largest clusters, respectively. **E)** Density plot showing the distribution of the fraction of phosphorylated conformers per protein, highlighting the prevalence of phosphorylation across the dataset.

To ensure reliable structural comparison, we curated a high-quality dataset by filtering for substantial fragments without missing residues. The final dataset includes 109 proteins with 253 phosphorylation sites, spanning all phosphorylation states-single, dual, and multiple-as well as corresponding non-phosphorylated structures (see Methods). Detailed information on the protein, structure and phosphorylation information is provided in Table S1.

### 3.1 Conformational diversity across phosphoproteins

To assess conformational diversity across phosphoproteins, structural conformers of each protein were clustered based on RMSD values (Figure 1A). Conformers are alternative structural states within a protein’s native ensemble that can range from large domain shifts to subtle local rearrangements, often arising from conditions such as substrate binding, post-translational modifications, or pH changes^22^. Here, we specifically define conformational diversity in the context of protein phosphorylation. We used the number of clusters per protein as a proxy for structural variability, where a higher cluster count indicates greater conformational diversity. As shown in the circular bar plot (Figure 1C), cluster counts varied across proteins, suggesting a broad range of structural flexibility in the dataset. Most proteins (66%) exhibited two distinct conformational states, followed by 26% with three states. Larger protein families, such as kinases, tended to show more clusters than smaller families like small GTPases and heat shock proteins.

To examine the composition of these clusters, we analysed the distribution of conformers within each. Figure 1D presents a dumbbell plot comparing the minimum and maximum cluster sizes for each protein, revealing differences in the representation of structural states within the PDB. Proteins with similar maximum and minimum conformer counts had balanced structural sampling, while those with wide disparities showed preferential representation of certain states-potentially phosphorylated forms-over others. This suggests a bias in structural deposition, favouring stable or experimentally tractable conformations.

We next explored whether phosphorylation contributes to this structural diversity. Almost 50% of proteins in our dataset contained at least 25% phosphorylated conformers (Figure 1E). To further investigate, we analysed the proportion of phosphorylated (P) and non-phosphorylated (NP) conformers within each cluster (Figure 2A). The heatmap shows clusters enriched for either P or NP conformers, as well as mixed clusters, indicating that phosphorylation can both influence and coexist within the structural landscape. Interestingly, 65% of proteins had at least half of their clusters containing both P and NP conformers (Figure 2A, inset). This overlap suggests that phosphorylation often stabilizes conformations already accessible in the NP ensemble, rather than inducing entirely new folds. In such cases, phosphorylation acts as a modulator-favouring specific pre-existing states rather than drastically altering structure-thereby enhancing the functional adaptability of the protein. Overall, these findings imply that the non-phosphorylated structural landscape is inherently flexible, encompassing conformations compatible with the phosphorylated state. Phosphorylation may therefore fine-tune protein function by shifting the equilibrium toward particular structural states in response to regulatory cues.

**Figure 2:**
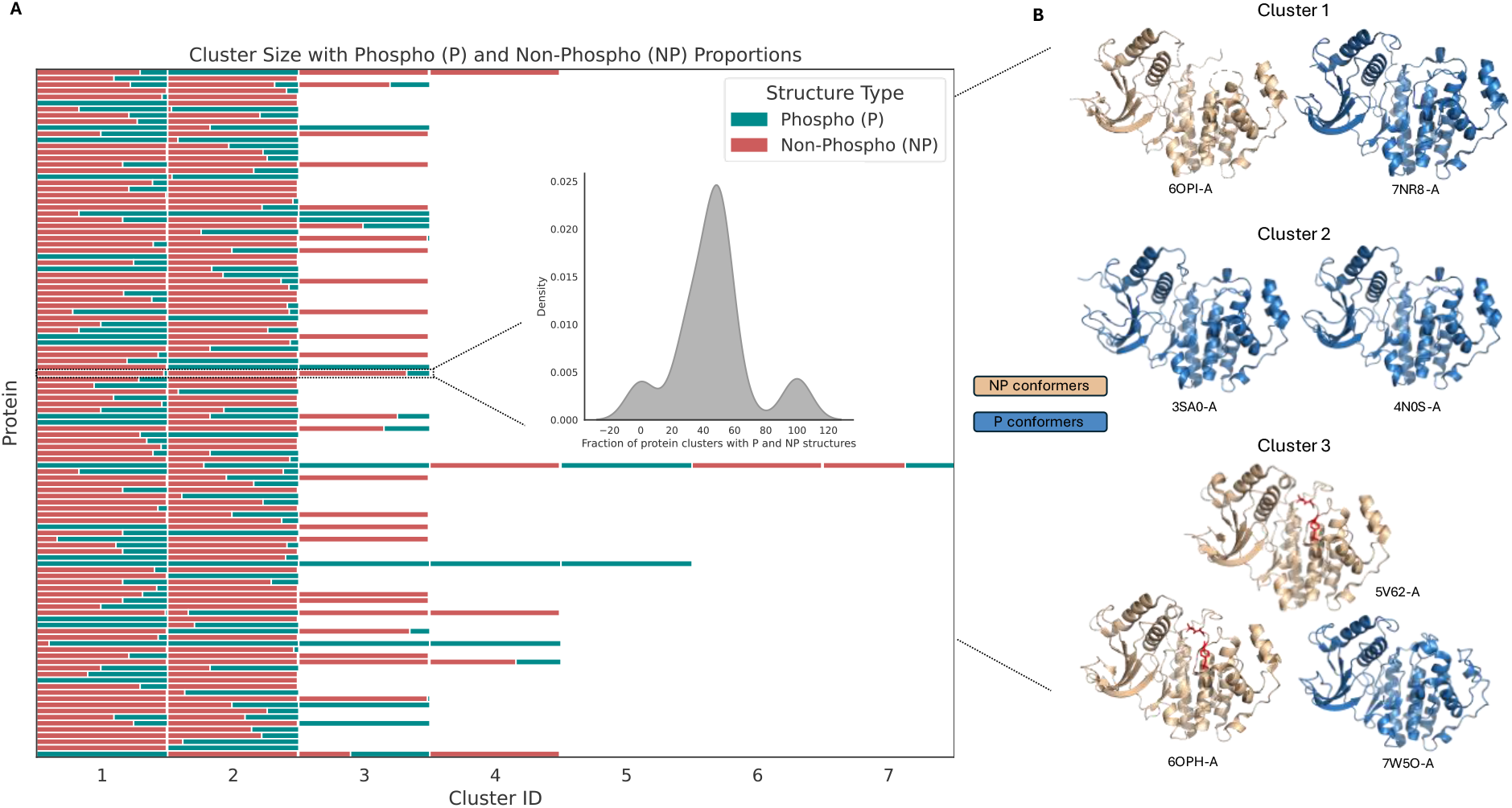
Mapping Conformational Diversity Across Phosphorylated Proteins. A) Heatmap illustrating the number of clusters (x-axis) for each protein (y-axis), with each cell representing a cluster. The proportion of P conformers is shown in cyan, and NP conformers in red. The inset highlights the fraction of clusters containing P and NP conformers across all proteins. B) Example from the dataset for Mitogen-activated protein kinase 1, which has three clusters. Cluster IDs 1 and 3 contain both P and NP conformers, while Cluster ID 2 consists exclusively of NP conformers. Representative structures for each cluster are shown, with P conformers in gold and NP conformers in blue. Phosphosites are highlighted in red and displayed as sticks where visible (e.g., in Cluster 1, the phosphorylated site in 6OPI-A is missing due to unresolved crystal structure regions).

### 3.2 Phosphorylation induced conformation shifts in experimental structures

To evaluate the structural impact of phosphorylation, we calculated pairwise RMSD values across three conformer pair types within each protein: P-P, NP-NP, and P-NP. RMSD values between P-P and NP-NP conformers were comparable, as shown by their proximity to the diagonal in Figure 3A, indicating similar levels of structural variability within each state. In contrast, P-NP comparisons exhibited higher RMSD values, clustering above the diagonal, suggesting that phosphorylation introduces greater structural divergence between P and NP states (Figure 3A).

**Figure 3:**
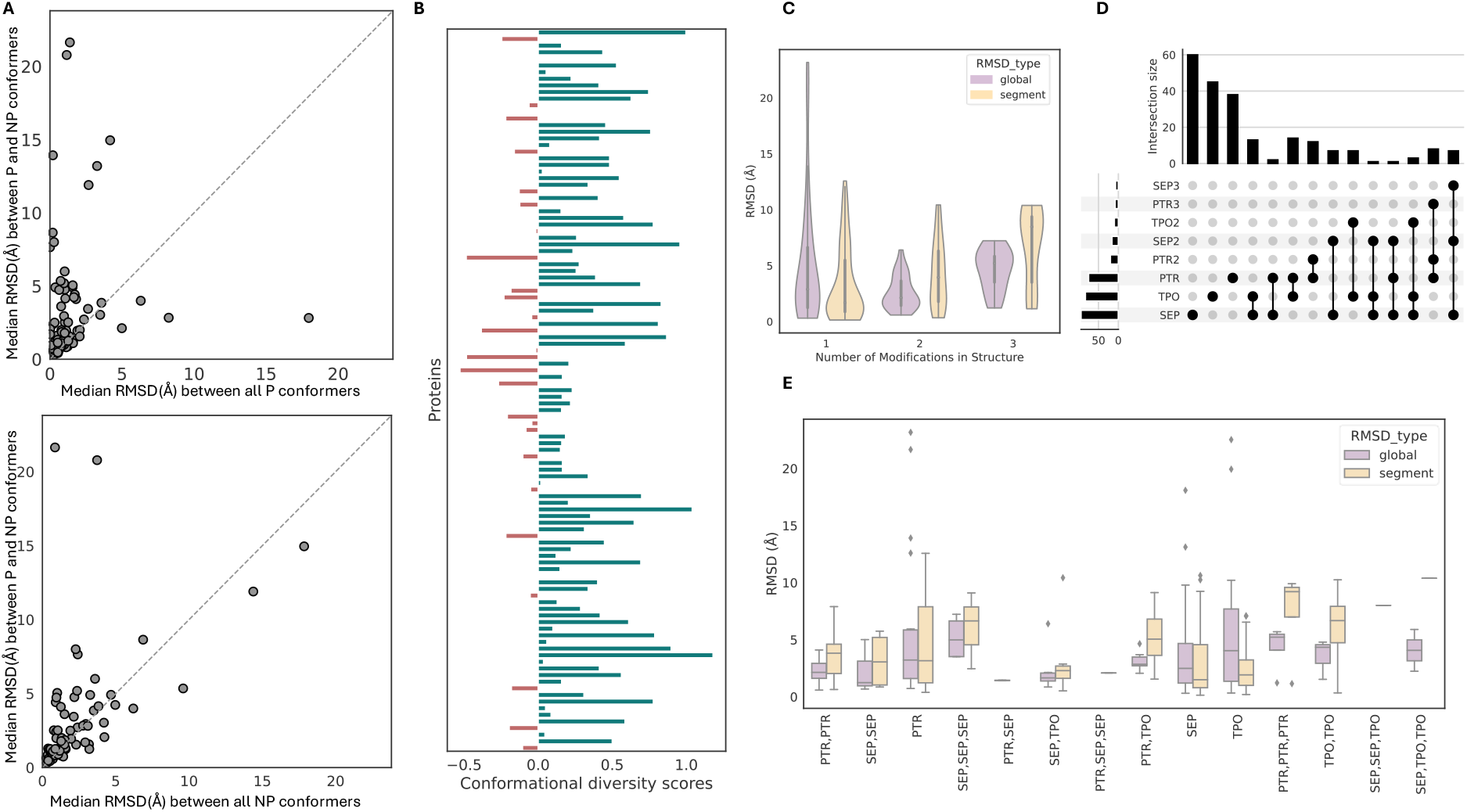
Conformational diversity and protein phosphorylation. A) Scatter plot (top) showing the median global RMSD values between P conformers (x-axis) and between P and NP conformers (y-axis) for each protein. Data points closer to the diagonal indicate less variability, while those further from the diagonal indicate higher diversity. The bottom scatter plot compares the median global RMSD between P and NP conformers (x-axis) and between all non-phospho conformers (y-axis) within each protein. B) Bar plot showing the difference between conformational diversity scores (CDS) and adjusted phospho CDS. Negative values indicates that phosphorylation induces greater conformational drift in those proteins. C) Residue-level modification analysis highlighting the most common single phosphorylation site (SEP), followed by TPO and PTR. Co-occurring phosphorylation sites are dominated by PTR-associated phosphosites. D) Violin plots showing the distribution of RMSD values for conformer pairs (P and NP) with the maximal RMSD in a protein. Global RMSD values are shown in purple, while segment RMSD values are in gold. The x-axis indicates the number of phosphosites in the phospho conformer. E) Boxplot showing RMSD values for the same dataset as in D, but with the x-axis representing the phosphosite combinations.

To assess whether molecular interactors contribute to phosphorylation-associated structural diversity, we compared RMSD distributions across P–P, P–NP, and NP–NP conformer pairs under different ligand-binding conditions. Ligand binding increased structural variability in NP conformers but had a stabilizing effect in P conformers. Correlation analyses further confirmed that phosphorylation was the primary driver of conformational divergence, while ligand effects acted as secondary modulators in a phosphorylation-dependent manner (see Supplementary Results and Figure S1 for detailed analysis).

#### 3.2.1 Conformational Diversity Score (CDS) and adjusted CDS Analysis

To quantify phosphorylation’s effect on structural diversity, we calculated the Conformational Diversity Score (CDS) and its phosphorylation-adjusted version (pCDS) for each protein (see Methods section 2.2.1 and 2.2.2). While CDS measures overall structural variability, pCDS normalizes this value by the proportion of P and NP conformers, isolating the contribution of phosphorylation. The difference between CDS and pCDS reflects phosphorylation’s structural impact. Positive values indicate that NP conformers drive most variability, with phosphorylation stabilizing pre-existing states (Figure 3B, S2A). Negative values suggest phosphorylation introduces novel conformations, expanding structural diversity (Figure 3B, S2C). Near-zero differences imply minimal structural shifts between P and NP conformers, pointing to a subtle regulatory effect (Figure 3B, S2B).

Overall, only 23% of proteins showed notable conformational drift upon phosphorylation, indicating that most proteins maintain similar structures across P and NP states under experimental conditions. Combined with the observed distributions of conformer fractions and median RMSD values (Figure 3A), these results suggest phosphorylation typically induces minor structural adjustments-stabilizing or subtly modulating NP conformations-rather than creating entirely new structural states.

#### 3.2.2 Conformational variability by phosphorylation state: global vs local effects

We evaluated how different phosphorylation states (single, dual, or multiple sites) affect structural variability by comparing phosphosites in experimental structures to their associated conformational shifts. For each protein, we selected the phospho state (single, dual, or multi) that exhibited the highest RMSD relative to the NP conformer. For instance, if a single modification (e.g., SEP) yielded higher RMSD than a dual-modified form (e.g., SEP+TPO), the single-modified conformer was used.

Conformers with a single phosphosite displayed the greatest global RMSD variability, with some showing extreme conformational shifts (~20 Å) (Figure 3C). This higher variability may reflect both the limited availability of multi-modified structures and the filtering strategy, where the phospho state showing the largest conformational shift relative to the NP state was selected (Figure 3C, 3D).Dual-modified structures exhibited narrower global RMSD distributions, suggesting a stabilizing effect, while triple-modified proteins showed a broader range and slightly elevated medians, indicating a possible interplay between stabilizing and destabilizing forces.

Segment-level RMSD remained tightly distributed across all modification states, indicating that conformational changes were localized-likely near the modification sites, possibly within intrinsically disordered regions (IDRs). Interestingly, while segment RMSD was lower than global RMSD in single-modified proteins, dual- and multi-modified states often showed higher segment-level median RMSD than their global values. This suggests that although global structures become more stable with increasing modification complexity, the local environment remains flexible and undergoes fine-tuned adjustments in response to multi-site phosphorylation.

#### 3.2.3 Conformational diversity and structural features

To understand the structural basis of phosphorylation-induced variability, we analysed domain organization, secondary structure transitions, and solvent accessibility in conformer pairs with maximal RMSD differences (as identified in Section 3.2.2). Proteins with phosphorylation sites located within structured domains exhibited greater conformational diversity, especially in multi-domain proteins. Phosphorylation-induced structural shifts were most prominent in flexible coil regions and were associated with increased solvent accessibility at the modification site. A detailed analysis is provided in Supplementary Results and Figures S3-S4.

### 3.3 Residue specific conformational change upon phosphorylation

Residue-specific domain analysis further confirmed the preference for modification sites, including both P and NP counterparts Tyrosine (TYR/PTR), Serine (SER/SEP), Threonine (THR/TPO), within structured regions. Modifications were predominantly observed in domain regions (PTR: n = 47; SEP: n = 41; TPO: n = 42), with fewer instances in non-domain regions (PTR: n = 6; SEP: n = 20; TPO: n = 2) (Figure S3C). Notably, SEP/SER displayed a higher occurrence in non-domain regions compared to PTR/TYR, TPO/THR. Mixed cases, where one conformer had a modification in a domain and the other in a non-domain region, were rare, with only one instance each for SEP/SER and TPO/THR and none for PTR/TYR (Figure S3C).

Our analysis suggests that sites corresponding to both P and NP states in conformers with maximal diversity pairs are predominantly localized in structured regions. However, SEP and SER exhibit greater flexibility, with a higher proportion occurring in non-domain regions compared to TYR, THR, and their P counterparts.

Residue-specific secondary structural analysis revealed that SER to SEP modifications most frequently maintained coil to coil states (n = 36) but were also observed in helix to helix states and occasionally transitioned from coil in the NP state to helix in the P state (Figure S4B). THR to TPO modifications exhibited transitions from helix in the NP state to coil in the P state, consistent with findings from a recent molecular dynamics study^39^, while PTR modifications were predominantly associated with strand states, either maintaining strand to strand transitions or adopting a strand conformation in the P state from a coil in the NP state. PTR and TPO modifications demonstrated larger changes in RMSD values and solvent accessibility compared to SEP, indicating a higher degree of structural reorganization in these residues (Figure S4C, D).

#### 3.3.1 Role of Single and multi-site phosphorylation in conformational diversity

We analysed global and segment RMSD values to assess residue-specific conformational variability driven by phosphorylation (Figure 3D). Segment RMSD was consistently higher than global RMSD, indicating increased local flexibility near modification sites, in line with previous observations of enhanced flexibility at phosphorylated residues^8^. Among single modifications, SEP showed the most stable conformations, with narrow RMSD distributions. TPO displayed intermediate variability, while PTR induced the largest segment-level shifts, reflecting pronounced localized reorganization (Figure 3E). In multi-site contexts, TPO-TPO and PTR-containing pairs (e.g., PTR-PTR, PTR-TPO) showed broader RMSD distributions and higher segment variability, whereas SEP-SEP pairs remained relatively stable. Overall, PTR drives the most substantial conformational changes, followed by TPO, while SEP is associated with minimal structural shifts (Figure 3E).

### 3.4 AlphaFold3 predicts the dominant structural state irrespective of phosphorylation status

AF’s ability to predict the effect of phosphorylation on proteins was tested on 109 protein pairs that showed little to large conformational differences upon phosphorylation. Most of these protein pairs are represented by kinase families, and most of these structural pairs are likely in the AF2 and AF3 training set. To evaluate the accuracy of AF2, AF3, and AF3-p predictions in capturing P and NP conformations, a systematic approach was undertaken. Pairwise RMSD values were calculated for all P vs. NP conformers, and the pair with the highest RMSD was selected to represent the most structurally diverse conformations for each protein.

This step ensured that the chosen conformers captured the maximum observed conformational variability between the two functional states. The predicted models (AF2, AF3, and AF3-p) were then aligned to these high-RMSD pairs to determine their proximity to either the P or NP conformational state. The RMSD values from this alignment were used to label each prediction as aligning with the P or NP state. This step provided an initial measure of how well the models captured the functional states associated with phosphorylation.

The results demonstrated that AF3-p predictions aligned with phosphorylated conformers in 73% of cases (80/109 proteins), slightly higher than AF3 (70%) and AF2 (64%) (Figure 4C). Our results suggest that AF3-p, which is tailored for phospho-specific predictions, exhibits a modest enrichment toward phosphorylated conformations compared to the general-purpose AF2 and AF3 models. However, the similarity in alignment proportions between the models indicates that AF2 and AF3 have comparable predictive preferences for P and NP states. Importantly, a chi-square test of independence (χ^2^=2.28, p=0.32) revealed no statistically significant differences in P/NP proportions across the models, indicating that all models perform similarly in aligning with P and NP conformers.

**Figure 4:**
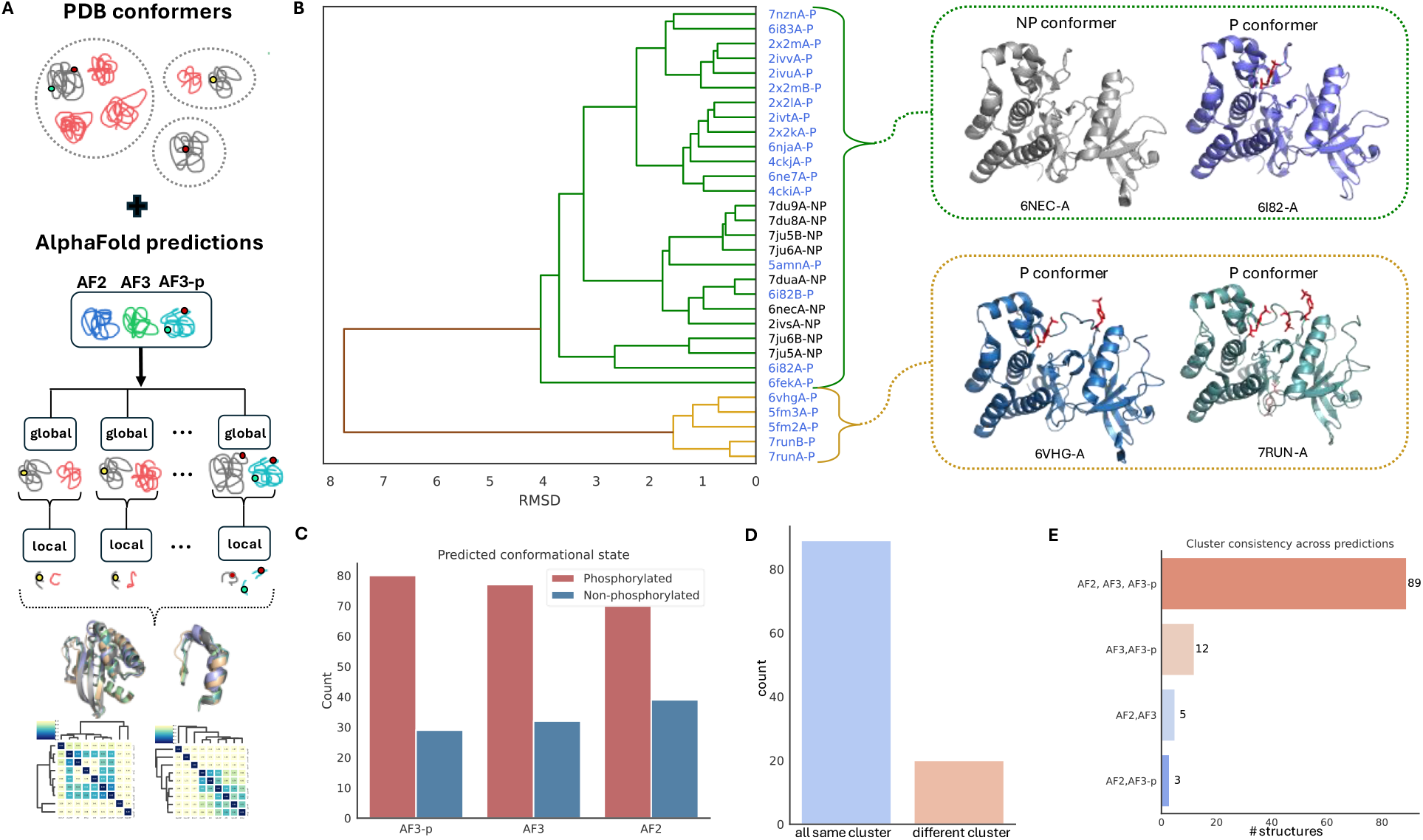
Evaluating AlphaFold predictions for phosphorylated and non-phosphorylated states. A) Workflow outlining the integration of PDB conformers with AlphaFold predictions (AF2, AF3, and AF3-p) for clustering based on global and local RMSD analyses. B) Hierarchical clustering of pairwise RMSD values for the Proto-oncogene tyrosine-protein kinase receptor (RET) reveals two distinct conformational clusters, suggesting the existence of two structural states. One cluster consists exclusively of P conformers, while the dominant cluster contains a mix of P and NP conformers. Representative structures from each cluster are displayed on the right, with phosphosites highlighted in red. C) Proportion of predicted conformational states aligning with the closest P or NP conformers for AF2, AF3, and AF3-p predictions. AF3-p shows a slight enrichment toward phosphorylated conformers compared to AF3 and AF2. However, all models exhibit a comparable tendency to predict P states over NP ones. D) Cluster consistency analysis reveals that 81.65% of AlphaFold predictions align with the same cluster across models, while 18.35% diverge into different clusters. E) Alignment of AlphaFold predictions with same dominant structural clusters indicates a strong preference for dominant conformational states across all models, regardless of phosphorylation specificity.

#### 3.4.1 Structural convergence and dominance in AlphaFold predictions

We evaluated the alignment of AlphaFold predictions (AF2, AF3, and AF3-p) to structural clusters corresponding to the closest P and NP conformational states. Hierarchical clustering of pairwise RMSD data grouped all conformers of a protein into structural clusters, with the cluster ID, size, and phosphorylation labels (See Methods, section 2.2.1 and 2.2.2) annotated for the conformer closest to each predicted model. This allowed us to assess whether the predictions from AF2, AF3, and AF3-p consistently aligned with the same cluster or spanned multiple distinct clusters. Across 109 proteins, we found that AlphaFold predictions aligned into the same cluster for 81.65% of cases, with predictions diverging into different clusters in only 18.35% of cases (Figure 4D, 4E). This high degree of alignment suggests a strong structural convergence across all three models, indicating that AlphaFold predictions are largely consistent with similar conformational states, regardless of phosphorylation status.

To further examine this tendency, we analyzed the alignment of predictions with dominant clusters, defined as the largest structural clusters for each protein. In cases where multiple clusters were of equal size, these clusters were categorized as “equal.” The results showed that all three models aligned with dominant clusters in 98% of cases. The Z-test for proportions revealed no significant differences in the proportion of dominant predictions among the three models (AF2, AF3, and AF3-p; p>0.05 for all pairwise comparisons), indicating that all models exhibit similar preferences for predicting dominant states, regardless of their specific design or focus on phosphorylation.

This consistent alignment with dominant clusters likely reflects patterns inherent to the training data, which emphasize the most stable or frequently observed conformations aligning well with the previous studies on fold switching proteins without modifications^27,28^. AlphaFold models thus appear more likely to favour dominant structural states rather than explore alternative or less-populated conformations, irrespective of phosphorylation specificity. This inherent bias highlights the model’s tendency to prioritize the most stable and frequently observed conformations, potentially limiting their ability especially AF3 to capture subtle structural variations and conformational shifts introduced by post-translational modifications such as phosphorylation. This pattern is further illustrated through a case study on the RET kinase receptor, where hierarchical clustering and heatmap analysis of RMSD values were used to evaluate the alignment of AlphaFold predictions with experimental conformers (see Supplementary Results and Figure S5).

#### 3.4.2 Assessing the structural fidelity of AlphaFold models across conformational ensembles

To address the potential bias of analyzing only a single pair of conformers, we expanded the analysis to include all available conformers for each protein that passed the filtering criteria (see Methods). This approach ensured that the structural diversity of both P and NP states was comprehensively represented, allowing for a more robust evaluation of AlphaFold model predictions against the full range of experimentally observed conformations. Pairwise RMSD calculations were performed between all conformers and the predicted models to identify the conformer closest to each prediction, with its phosphorylation status recorded. This expanded alignment offered a more comprehensive evaluation of how effectively the models captured the full range of structural diversity observed in the PDB, rather than focusing solely on a single pair exhibiting high conformational variability.

For example, for each model type (AF2, AF3, AF3-p), we first identified the closest experimental conformer (P or NP) from the pair showing the largest conformational difference for each protein. We then retrieved the corresponding cluster ID, cluster size, and phosphorylation state (“P” or “NP”) for that conformer. To expand beyond this single high-variability pair, we included all available P and NP conformers, along with the predicted models, in hierarchical clustering. From the resulting dendrogram, we again identified the closest experimental conformer for each model type and extracted its cluster information, including ID, size, and phosphorylation state.

We then compared the closest conformers identified from the high-RMSD pair with those identified through comprehensive clustering. If both the cluster ID and phosphorylation state matched, the prediction was considered to have captured the same conformational and functional state. If the cluster ID matched but the phosphorylation state differed, it suggested that the model predicted a similar structural state, independent of phosphorylation status. We also examined the cluster size of the closest conformer to assess whether the models aligned with the most dominant structural state.

Predictions were categorized based on whether they aligned with the “same” or “different” clusters, and whether the phosphorylation states matched or differed. This allowed us to evaluate the AlphaFold model’s ability to reproduce both structural similarity and phosphorylation-dependent conformational differences. The analysis quantified how often models aligned with similar structural clusters, and to what extent they sampled structurally diverse or functionally distinct states.

Across all AlphaFold models (AF2, AF3, AF3-p), predictions predominantly aligned with the same cluster as the closest experimental conformer. AF3-p showed the highest cluster alignment (81%), followed by AF3 (80%) and AF2 (71%). A Kruskal-Wallis test based on binary classification (“same” vs. “different” cluster) revealed no significant differences among the models (H = 4.20, p = 0.12), indicating comparable performance in structural cluster matching across AF2, AF3, and AF3-p.

When phosphorylation state was considered, predictions that aligned with the same structural cluster generally also matched in phospho-state. AF3 had the highest number of matches (n = 50), followed by AF3-p (n = 49) and AF2 (n = 43). In contrast, predictions aligning with different clusters were more likely to diverge in phosphorylation state, with AF2 showing the highest number of mismatches (n = 21).

Binomial tests assessing enrichment for phospho-state matches revealed that none of the models significantly favored matching phosphorylation states beyond random expectation (AF3-p: p = 0.44; AF3: p = 0.70; AF2: p = 0.84). Similarly, a chi-squared test comparing the overall distribution of matching vs. non-matching phospho-states across models showed no significant difference (χ^2^ = 0.68, p = 0.71). Overall, while AF3-p showed a slight trend toward better phospho-state agreement, all models performed similarly in both structural cluster alignment and phosphorylation state matching. These findings highlight the need for further optimization to improve model sensitivity to phosphorylation-dependent conformational changes.

#### 3.4.3 Co-evolutionary constraints drive AlphaFold model bias toward dominant protein conformations

To investigate why AF models favour specific conformational states, we examined structural and evolutionary interaction patterns, focusing on global residue interaction networks, localized contacts around P and NP sites, and co-evolutionary relationships inferred from co-evolving residue pairs. We selected three proteins with distinct conformational landscapes: P04406, where P and NP states are nearly identical; P24842, where NP states drive conformational variability while P states remain stable; and P07949, which exhibits multiple P conformers, with one driving structural diversity while others coexist with NP states.

Across all cases, we observed that AlphaFold predictions favoured evolutionarily stable conformations, those reinforced by co-evolutionary constraints rather than sampling the full spectrum of conformational variability (Figure 5). AF3-p consistently aligned with dominant P states that coexist with NP conformers, while AF2 and AF3 more frequently predicted NP conformers, especially in cases where NP states were responsible for the observed conformational diversity.

**Figure 5:**
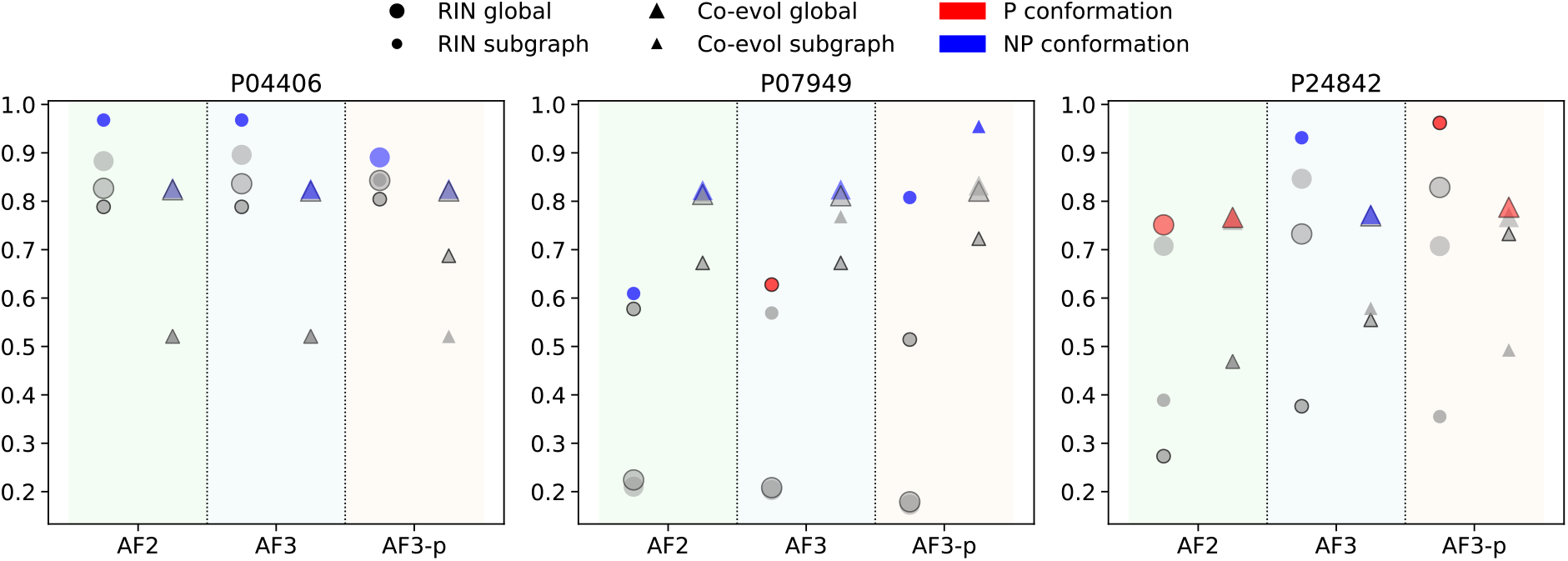
Comparison of predicted models to experimental PDB conformers based on similarity scores derived from global residue interaction networks, subgraph-level interactions, and co-evolving residue pairs. The x-axis denotes the type of predicted model, while the y-axis indicates the similarity score relative to the PDB conformer. The highest scores in each category are highlighted, with respect to the phosphorylation status of the PDB conformer.

In P04406, where P and NP conformers are structurally indistinguishable, all three models accurately predicted the NP state (Figure 5). This suggests that in the absence of conformational diversity, AlphaFold models generalize well and reproduce both P and NP states with high fidelity. In P07949, AF2 and AF3 predicted a structurally distinct P conformer that drives variability, whereas AF3-p aligned with a coexisting P conformer that clusters closely with NP states (Figure 5). This indicates that AF3-p does not simply favour phosphorylation but is instead tuned to detect P states that are evolutionarily constrained and structurally compatible with dominant NP conformations.

Similarly, in P24842, AF2 and AF3 aligned with NP conformers-AF3 in particular with the NP state that drives diversity-while AF3-p favoured a P/NP coexisting conformer that resides in a more evolutionarily stable cluster (Figure 5). These findings are reinforced by higher co-evolutionary subgraph similarity scores for AF3-p in dominant clusters, highlighting that AF3-p is better optimized to capture phosphorylated conformations that are embedded within conserved structural environments (Figure 5).

Together, our findings suggest that AF models, rather than learning all viable conformations equally, are optimized to reproduce the most frequently observed or evolutionarily stable states. AF3-p’s preference for dominant P conformers suggests that phosphorylated residues within stable environments exert a strong influence on model predictions, whereas AF2 and AF3 remain more sensitive to NP conformers, particularly when these dominate the conformational landscape.

## 4 Discussion

Our study provides an integrated perspective on how phosphorylation modulates protein structure and evaluates the ability of AlphaFold models (AF2, AF3, and AF3-p) to capture these changes. By combining a systematic analysis of experimentally resolved phosphorylated (P) and non-phosphorylated (NP) structures with an assessment of AlphaFold predictions, we uncover critical insights into conformational landscape, model biases, and implications for the broader research community.

### 4.1 Phosphorylation as a modulator of structural shifts

Phosphorylation predominantly stabilizes or subtly adjusts pre-existing structural states rather than inducing novel conformations^6^. Approximately 65% of P and NP conformers share the same structural cluster, suggesting that the NP ensemble often includes conformations accessible to the P state. These findings highlight phosphorylation’s role as a fine-tuner of protein function, favouring specific structural states that support regulatory or signalling processes. However, distinct conformational shifts upon phosphorylation were observed in 23% of proteins, underscoring the context-dependent nature of these modifications. These results challenge the notion of phosphorylation as a driver of dramatic structural transitions and instead position it as a modulator of existing flexibility, which aligns with the previous study^6^.

Our results reveal that ligand binding modulates conformational diversity differently depending on phosphorylation status. Ligand binding reduced diversity in P conformer pairs (P-P, P-NP) but increased it in NP pairs (NP-NP), suggesting that phosphorylation constrains structural flexibility while NP states remain more ligand-responsive. Weak correlations between P-P and NP-NP pairs indicate that phosphorylation primarily drives structural divergence, while strong correlations in P-NP vs. NP-NP pairs suggest ligand binding amplifies structural overlap in mixed phosphorylation states. These findings highlight phosphorylation as the primary determinant of conformational diversity, with ligand binding playing a secondary, context-dependent role.

Residue-specific analyses reveal that structures with single phosphorylation sites exhibit the highest global conformational variability, whereas multi-site modifications tend to balance localized flexibility with overall structural stability, highlighting a nuanced interplay between stabilization and disruption. Phosphotyrosine (PTR) and phosphothreonine (TPO) induce the most pronounced structural shifts, both globally and locally, reflecting their dynamic roles in functional regulation. As a single modification, TPO drives greater global structural changes, while phosphoserine (SEP) shows intermediate variability and minimal impact, aligning with its role in stabilization. Notably, dual modifications involving PTR and TPO, either as identical pairs (e.g., TPO-TPO or PTR-PTR) or combinations (e.g., PTR-TPO), lead to pronounced shifts, particularly in localized protein conformations.

### 4.2 Biases in experimental data and their impact on predictions

The PDB data underpinning AlphaFold’s training exhibits pronounced biases, with phosphorylated structures disproportionately represented in kinase superfamilies and limited diversity in multi-site modifications. This imbalance might skew AlphaFold predictions toward dominant, well-represented conformational states, at the expense of rare or functionally significant phosphorylation-induced structures. The overlap between P and NP conformers within the same cluster further reflects the constrained diversity of experimental data, limiting the ability of models to capture the broader functional implications of phosphorylation. These findings emphasize the need for more diverse datasets to enhance the fidelity of predictive models. Our study also highlights the essential dearth of such data, severely limiting model training and validation.

### 4.3 AlphaFold performance: structural memorization and functional fidelity

Despite the inclusion of PDB-derived structural information in their training, AF2, AF3, and AF3-p struggle to differentiate between P and NP conformers effectively. Our results show that while AF3-p demonstrates a slight enrichment toward phosphorylated states, the differences in predictive performance across models are not statistically significant. All models exhibit a strong bias toward dominant conformational states, aligning with the largest clusters irrespective of phosphorylation status. This tendency reflects a structural memorization effect, where AF models prioritize frequently observed conformations in the training data rather than capturing phosphorylation-induced structural diversity^27,28^.

Furthermore, AF3-p’s inability to substantially outperform AF2 and AF3 raises questions about its phospho-specific design. While AF3-p aligns more closely with phosphorylated conformers in some cases, these instances remain exceptions rather than the rule. This limitation underscores the challenge of overcoming biases in training data and highlights the need for model refinement to improve sensitivity to phosphorylation-specific structural changes.

### 4.4 Implications for the research community

The biases and limitations identified in AlphaFold predictions have significant implications for researchers studying post-translational modifications (PTMs). While AlphaFold models excel at predicting dominant conformations, their inability to capture rare or biologically significant phosphorylation-induced transitions necessitates cautious interpretation. These findings reinforce the importance of integrating computational predictions with experimental validation, particularly for dynamic modifications like phosphorylation.

Addressing the limitations identified in this study requires a two-pronged approach: improving training datasets and refining model architectures. Expanding structural datasets to include more phosphorylated protein structures with diverse modification states will provide the foundation for better model training. In parallel, enhancing model architectures to prioritize local structural changes and rare conformations over dominant ensembles may improve their ability to predict phosphorylation-induced diversity.

Additionally, integrating advanced techniques such as transfer learning, where models can adapt to phosphorylation-specific tasks using high quality, curated datasets, may enhance their performance. Complementary approaches, such as combining AF predictions with molecular dynamics simulations and experimental validation, could further bridge the gap between predicted and experimentally observed conformations.

Refining AlphaFold architectures to prioritize local structural changes and rare conformations over dominant ensembles represents a promising avenue for improvement. Incorporating advanced techniques such as context-specific embeddings or hybrid predictive frameworks may enable more accurate modeling of phosphorylation-induced structural landscapes.

In conclusion, phosphorylation acts as a nuanced regulator of protein structure, favoring functional adaptability through subtle modifications rather than dramatic shifts. While AlphaFold models remain invaluable tools for structural biology, their biases toward well-sampled states limit their applicability to phosphorylation-specific studies. By addressing these challenges, the research community can drive the development of more accurate and functionally relevant predictions, unlocking deeper insights into the dynamic roles of PTMs in cellular regulation and disease.

## Supporting information

Supplementary Results

## Data availability

All data analyses and plots were done in Python (version 3). Protein structure figures were generated using PyMol^40^ (version 3.0.5). All of the data and python scripts used in these methods are available online (https://github.com/Pathmanaban/AlphaFold-phospho-conformation).

## 5 Acknowledgment

PR acknowledges BOF grant from Ghent University under grant agreement number BOF.PDO.2023.0024.01. PR, JZ, LM and WV acknowledges Research Foundation Flanders (FWO) through grant G028821N.

