## Supplementary Results for "Assessing the relation between protein phosphorylation, AlphaFold3 models and conformational variability"

### The influence of molecular interactors on phosphorylated and non-phosphorylated conformers

We further examined whether molecular interactors-such as ligands, ions, or peptide binders-modulate conformational diversity differently in P and NP conformers. RMSD distributions were compared across all pair types (P-P, P-NP, NP-NP) under three ligand-binding conditions: both conformers ligand-bound, only one conformer ligand-bound (mixed), and neither conformer ligand-bound (no-ligand) (Figure S1).

Tukey HSD tests showed that ligand binding significantly affected RMSD distributions. In NP-NP pairs, RMSD values were highest when both conformers were ligand-bound, indicating increased conformational variability. In contrast, ligand binding reduced RMSD in both P-P and P-NP pairs, suggesting a stabilizing effect in phosphorylated conformers.

Correlation analysis further clarified the differential influence of ligand binding on structural diversity. Weak or non-significant correlations were observed between P-P and P-NP pairs across ligand-binding conditions ($r=0.23$, $p=0.10$) and in no-ligand conditions ($r=0.26$, $p=0.03$), indicating that phosphorylation induces conformational shifts largely independent of ligand effects. Similarly, P-P vs. NP-NP comparisons yielded only weak to moderate correlations ($r=0.41$ for ligand-bound pairs, $r=0.27$ for no-ligand pairs) reinforcing phosphorylation’s role as the main driver of conformational differences.

In contrast, strong correlations were observed between P-NP and NP-NP pairs, especially in ligand-bound ($r = 0.74$, $p < 0.001$) and mixed conditions ($r = 0.79$, $p < 0.001$), suggesting that ligand binding strongly influences NP conformers but has a limited, stabilizing role in P conformers. Scatter plots (Figure S1) illustrate these trends: NP-NP pairs showed broader RMSD variability when ligand-bound, while P-P and P–NP pairs remained more structurally constrained. Our findings indicate that ligand binding enhances structural variability in NP conformers but stabilizes P conformers. Phosphorylation emerges as the primary determinant of conformational divergence, with ligand effects acting as secondary modulators in a phosphorylation-dependent manner.

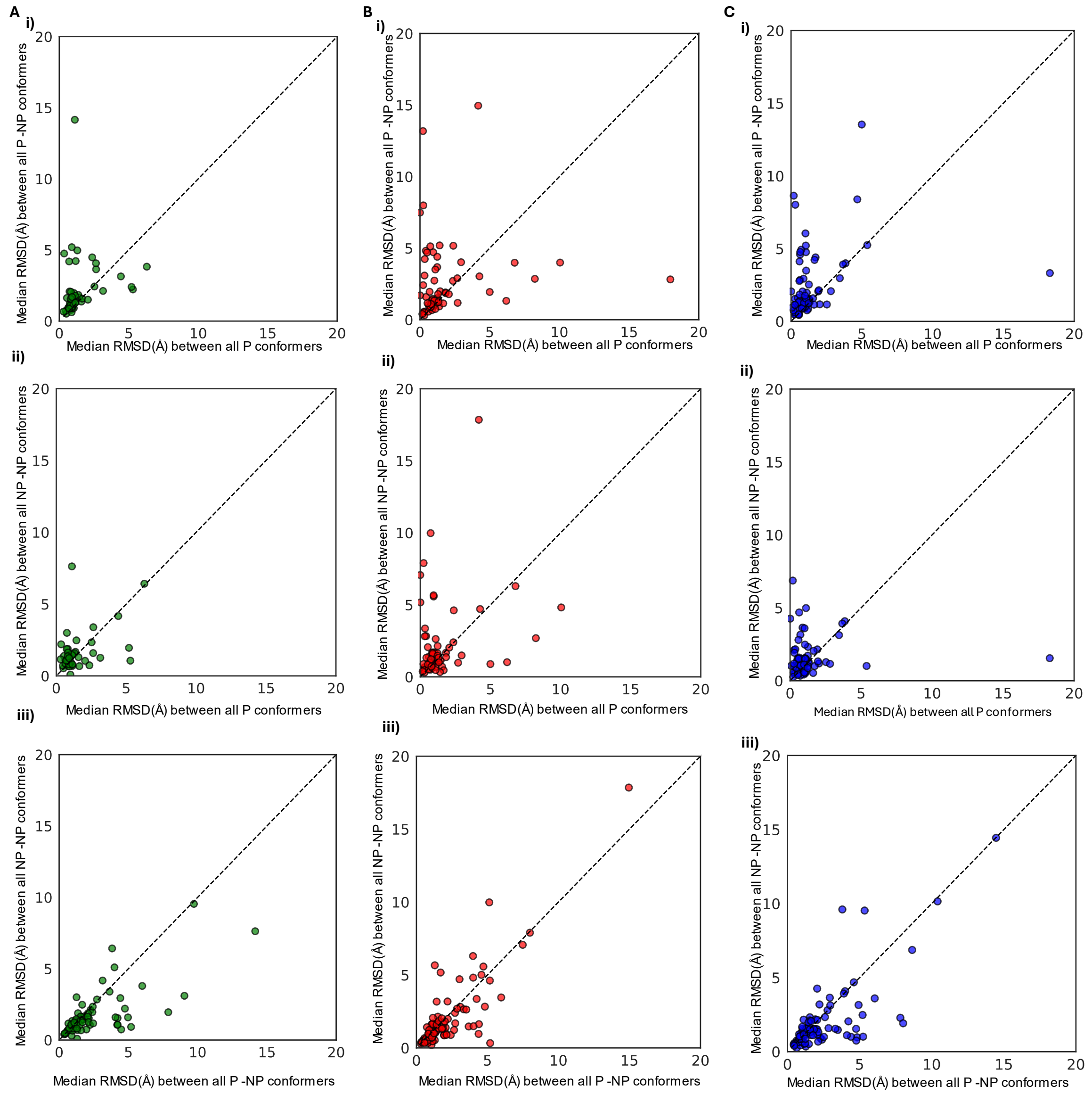


**Figure S1: Conformational diversity, molecular interactors, and phosphorylation.** A) Scatter plot (left) showing the median global RMSD values between P and NP conformers in ligand-bound states. B) Scatter plot (middle) illustrating the median global RMSD values between P and NP conformers in no-ligand states. C) Scatter plot (right) depicting the median global RMSD values for mixed-ligand states, where one conformer (P or NP) is ligand-bound while the other is not. i) Median RMSD between all P conformers (x-axis) and P-NP conformers (y-axis). ii) Median RMSD between all P conformers (x-axis) and all NP conformers (y-axis). iii) Median RMSD between all P-NP conformers (x-axis) and NP conformers (y-axis). Data points near the diagonal reflect lower variability, while those farther from the diagonal indicate greater structural diversity.

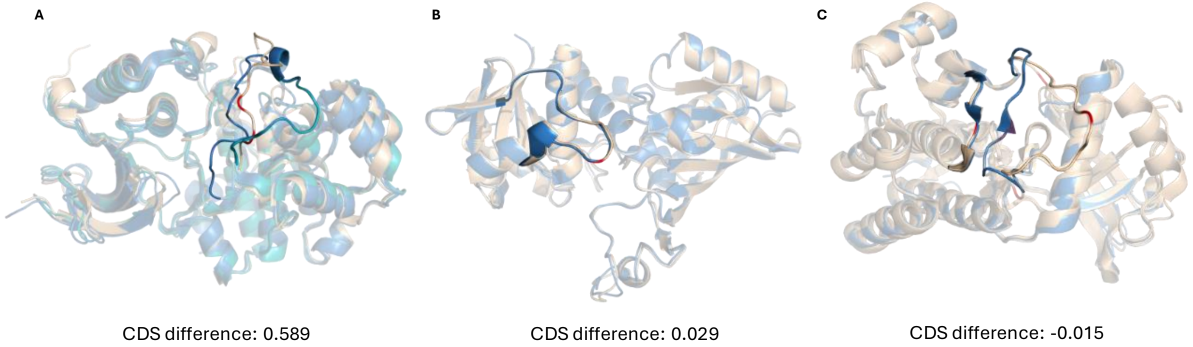


**Figure S2: Structural adaptations of phosphorylated protein conformers.** Examples of protein conformational landscapes observed in experimental structures and their conformational diversity score (CDS) differences. A) The CDS difference for Mitogen-activated protein kinase 1 (MPAK1) shows a positive value (0.589), indicating that NP conformers primarily drive the conformational diversity, while phosphorylation stabilizes the pre-existing NP conformation. Structural alignment of P (in gold, phosphosites in red) and NP conformers (in blue and cyan) shows that the NP state exists in multiple conformational ensembles, with the P conformer being part of these pre-existing states. B) Glyceraldehyde-3-phosphate dehydrogenase (GADPH) has a CDS difference near zero (0.029), indicating no significant difference between P and NP conformers. Structural alignment shows strong similarity between P (in gold, phosphosites in red) and NP states (in blue). C) Proto-oncogene tyrosine-protein kinase receptor Ret (RET) has a negative CDS difference (-0.015), suggesting that phosphorylation drives conformational diversity. Alignment of P (in gold) and NP conformers (in blue) shows certain phospho states that exist in a distinct conformational state different from the NP state. For clarity, only selected P and NP conformers are shown in all examples, with the fragments of interest highlighted.

### Conformational diversity and structural features: domain architecture, secondary structure, and solvent accessibility

To investigate the relationship between domain organization and conformational diversity between P and NP pairs with maximal diversity (i.e., pairs exhibiting the highest RMSD values as identified in Section 3.2.2), RMSD values and domain annotations were compared across single-domain (when both P and NP conformers are modelled with a single domain), multi-domain (when the conformers are modelled with more than one domain with possible linkers), and mixed-domain proteins (where one conformer is modelled as a single domain and the other as multi-domain).

After annotating the domain organization for the maximal diversity pairs, we found that the conformers were predominantly composed of single-domain proteins (n=73), followed by multi-domain proteins (n=22) and mixed-domain pairs (n=9) (Figure S3A,B). RMSD values were generally more consistent in single-domain proteins, while multi-domain proteins exhibited greater structural divergence, reflected by a wider spread of RMSD values (Figure S3A). Although the spread between single and multi-domain proteins was not substantial, extreme cases of structural variability were observed in multi-domain proteins. Mixed-domain proteins showed intermediate RMSD values but were underrepresented in the dataset (Figure S3A).

Analysis of domain localization at phosphosites revealed that the majority of modifications (n=130) had both modified and unmodified sites located within structured domain regions, while a smaller subset (n=28) had both sites outside domain regions, such as in unstructured or linker regions. Only two sites were observed in the mixed category, where one conformer had a modified residue in a domain while the other was located outside a domain (Figure S3B). These findings highlight that maximal conformational diversity emerges when modifications are positioned within structured regions of the protein, hinting at a deeper interplay between structural organization, secondary structure transitions, and solvent exposure.

Differences in secondary structure and solvent accessibility between P and NP conformers were analysed using STRIDE annotations, consolidating eight structural classes into three categories: coil, helix, and strand. State changes and solvent accessibility variations were compared between NP and P conformers (Figure S4). The majority of secondary structure states remained stable, with coil to coil (n=91) being the most prevalent, followed by strand to strand (n=18) and helix to helix (n=15). RMSD analysis revealed greater variability in flexible regions, particularly in transitions involving coil to coil, helix to coil, and coil to helix states (Figure S4A). Structured regions, such as helix to helix and strand to strand, showed as expected relatively lower RMSD variability (Figure S4C). Additionally, transitions such as coil to strand and strand to coil exhibited consistent RMSD values. Changes in solvent accessibility indicated an overall increase in accessibility upon phosphorylation (Figure S4D).

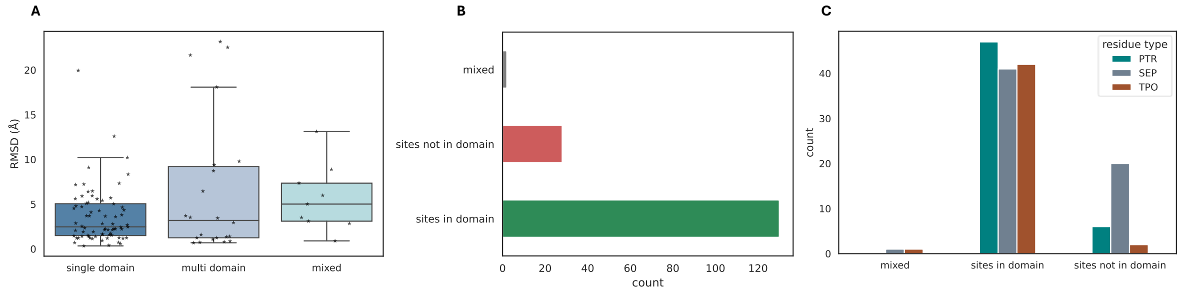


**Figure S3: Domain architecture and protein phosphorylation.** A) Distribution of RMSD values between proteins with single-domain conformers (in both P and NP pairs, n=73), multi-domain conformers (in both P and NP pairs, n=22), and mixed-domain conformers (where one pair is single-domain and the other is multi-domain, n=9). B) Plot showing the distribution of P and NP sites located in domain and non-domain regions for maximal diversity pairs. "Domain" refers to cases where both P and NP sites in the pair are located within domain regions, "not in domain" refers to cases where both P and NP sites are located outside domain regions, and "mixed" refers to cases where one site (P or NP) is located within a domain and the other is not. C) Same data as B, but presented at the residue-specific level.

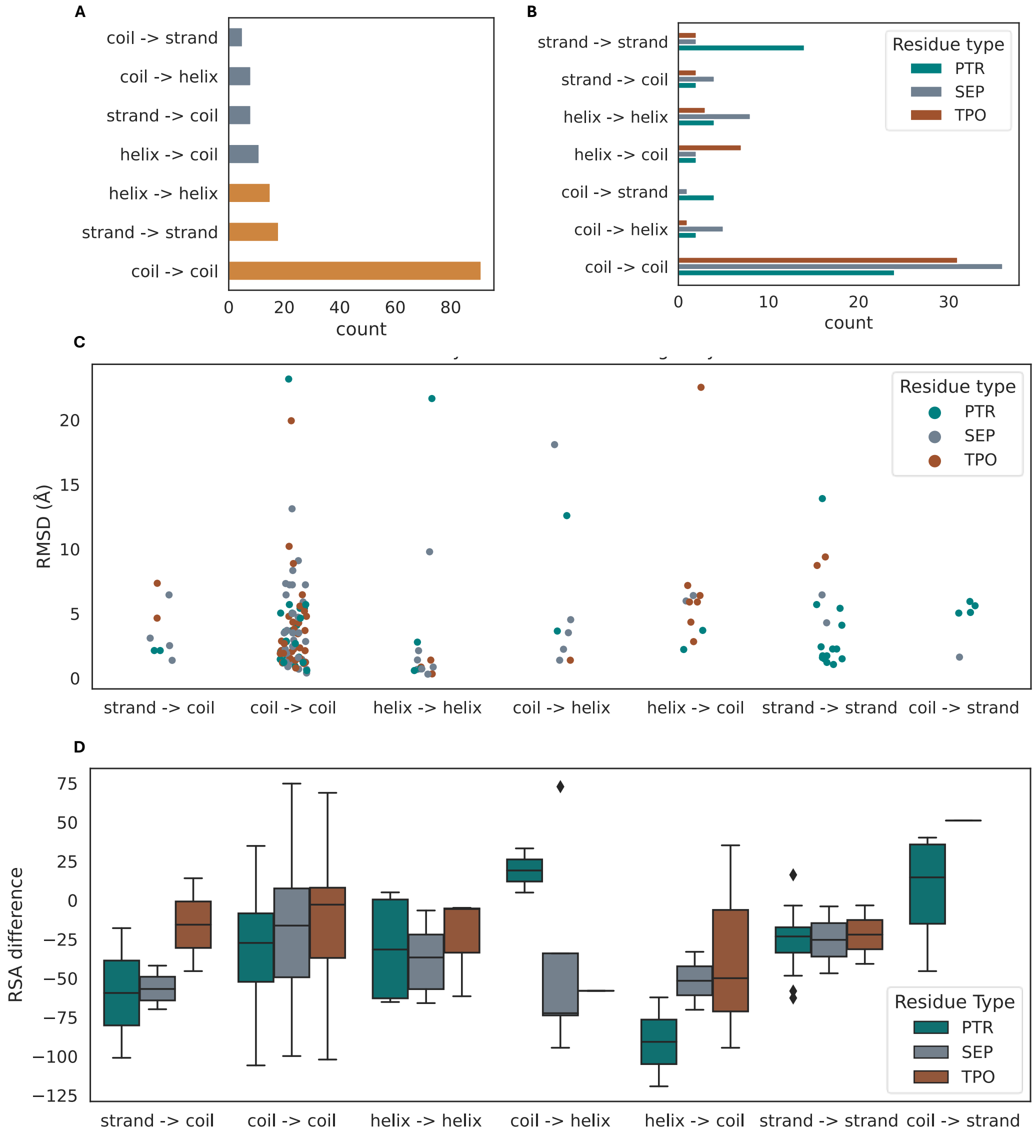


**Figure S4: Secondary structural state transitions and solvent accessibility changes upon phosphorylation.** A) Distribution of secondary structural state changes from NP to P conformers observed in maximal diversity pairs. B) Residue-specific secondary structural state changes in maximal diversity pairs. C) Distribution of residue-specific secondary structural changes (x-axis) from NP to P states and their corresponding RMSD values (y-axis). D) Differences in relative solvent accessibility of residues from NP to P states, with negative values indicating an increase in solvent accessibility upon phosphorylation.

### Assessing local conformational changes through segment and contact analysis

To complement the dominance and convergence analyses, we further examined global and local conformational changes upon phosphorylation to visually assess the alignment of AlphaFold predictions (AF2, AF3, and AF3-p) with experimental structures. Since global RMSD values often overestimate local structural shifts, segment alignments were performed using 15 residues flanking the phosphorylation site (N- and C-termini). Lower segment RMSD values between AlphaFold models and conformers indicated closer alignment with experimentally observed conformations.

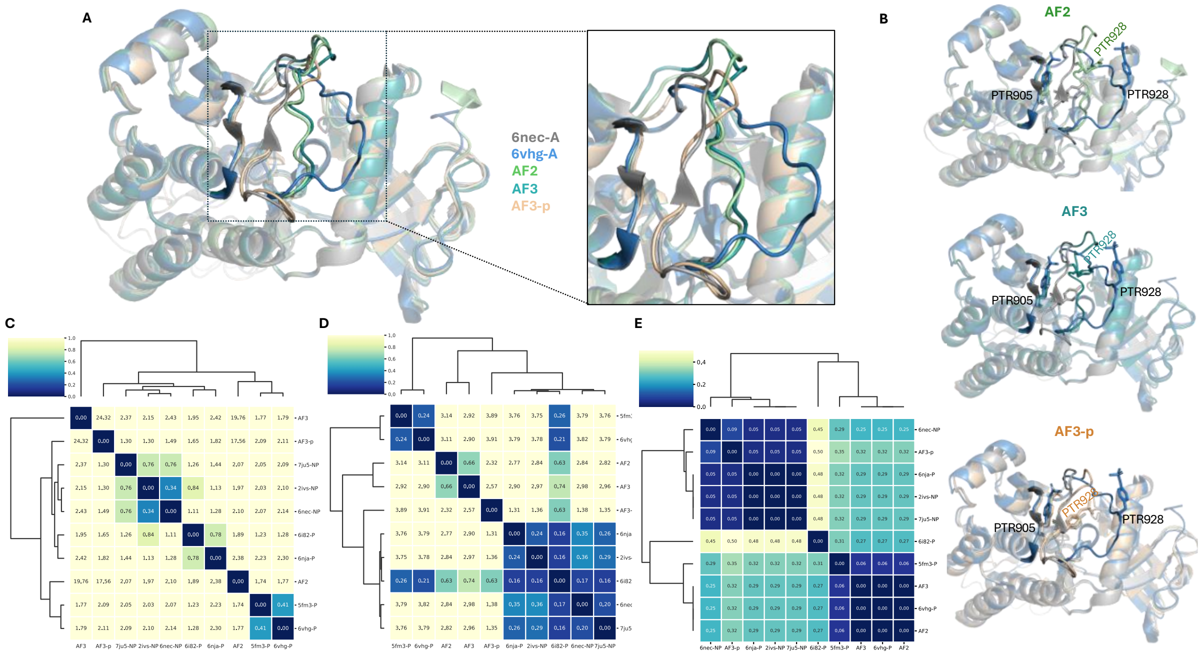


**Figure S5: Conformer alignment and hierarchical clustering of AlphaFold predictions with P and NP conformers** A) Structural alignment of the Proto-oncogene tyrosine-protein kinase receptor (RET) (UniProt ID: P07949), showing P and NP conformers alongside AlphaFold predictions (AF2, AF3, and AF3-p). The NP conformer (grey) and the P conformer (blue), which exhibited the maximum RMSD for the protein, are aligned with AF2 (green), AF3 (dark cyan), and AF3-p (gold) predictions. The inset highlights the phosphorylation segment and its alignment with the predicted models. B) Individual alignments of AF2, AF3, and AF3-p predictions with P and NP conformers, illustrating differences in structural alignment across the models. Phosphosites are depicted as sticks. C) Heatmap of global RMSD values between experimental P and NP conformers and AlphaFold predictions, hierarchically clustered to group structurally similar conformations. D) Heatmap of segment RMSD values (15 residues flanking the phosphorylation site), capturing local structural shifts between experimental conformers and predictions. E) Heatmap of contact map differences (within an 8 Å radius of phosphosites), hierarchically clustered to assess alignment of AlphaFold predictions with residue-level contact environments of P and NP conformers. Across all heatmaps, AF3-p predictions consistently aligned with the dominant cluster (Figure 4B dendrogram), even when these clusters included members displaying conformational states distinct from the modeled phosphosites (PTR 905 and PTR 928).

Condensed distance matrices were generated using global RMSD, segment RMSD, and contact map differences (residues within 8 Å of P/NP sites). For each phosphorylation state, the best representative conformers were selected along with the NP conformers to construct these matrices, and AlphaFold predictions were clustered alongside these conformers to evaluate alignment with structural ensembles.

Clustering revealed distinct patterns in the alignment of AlphaFold predictions with P and NP conformers. While global RMSD analysis often failed to detect subtle conformational differences around phosphorylated regions, segment-level RMSD and residue contact map analyses were more effective in capturing these minor variations. For example, the Proto-oncogene tyrosine-protein kinase receptor (RET) exists in multiple phosphorylation states, and hierarchical clustering identified two distinct conformational clusters for this protein (Figure 4B), corresponding to two conformational states (Cluster 1 and Cluster 2). The P conformer (PDB ID: 6VHG, chain A), which exhibited the largest conformational shift to the NP state for this protein, contained two phosphosites (PTR 905 and PTR 928).

For example, for protein **Proto-oncogene tyrosine-protein kinase receptor (RET),** heatmap of global RMSD, segment RMSD and contact map clustering (Figures S5C, S5D, S5E) showed that AF3-p aligned with the most dominant cluster irrespective of phosphorylation status, while AF2 and AF3 showed varying cluster alignment (Figure S5D, S5E). Although this discrepancy in cluster assignment was observed for 18.4% of cases, the majority of proteins (81.6%) showed all three models aligning with the most dominant conformational state. These results highlight the structural biases of the models, with the phosphorylation-agnostic AF2 and AF3 favoring well-represented conformations. Interestingly, AF3-p's phospho-specific training did not result in significantly distinct predictive behavior, as it predominantly also aligned with dominant clusters in a manner similar to AF2 and AF3.

Therefore, while AF3-p at first sight appears to perform better for phosphorylated states, a closer examination shows that AF2 and AF3 align with P conformers at rates comparable to AF3-p. This convergence across models suggests that AF3-p's performance is not substantially distinct, reinforcing the observation that all models are heavily influenced by structural patterns and dominant conformational ensembles present in the training data. This bias toward well-sampled states, irrespective of phosphorylation status, underscores a tendency for structural memorization rather than capturing the functional landscape associated with phosphorylation.
